# Evidence for a mouse origin of the SARS-CoV-2 Omicron variant

**DOI:** 10.1101/2021.12.14.472632

**Authors:** Changshuo Wei, Ke-Jia Shan, Weiguang Wang, Shuya Zhang, Qing Huan, Wenfeng Qian

## Abstract

The rapid accumulation of mutations in the SARS-CoV-2 Omicron variant that enabled its outbreak raises questions as to whether its proximal origin occurred in humans or another mammalian host. Here, we identified 45 point mutations that Omicron acquired since divergence from the B.1.1 lineage. We found that the Omicron spike protein sequence was subjected to stronger positive selection than that of any reported SARS-CoV-2 variants known to evolve persistently in human hosts, suggesting the possibility of host-jumping. The molecular spectrum (*i*.*e*., the relative frequency of the twelve types of base substitutions) of mutations acquired by the progenitor of Omicron was significantly different from the spectrum for viruses that evolved in human patients, but was highly consistent with spectra associated with evolution in a mouse cellular environment. Furthermore, mutations in the Omicron spike protein significantly overlapped with SARS-CoV-2 mutations known to promote adaptation to mouse hosts, particularly through enhanced spike protein binding affinity for the mouse cell entry receptor. Collectively, our results suggest that the progenitor of Omicron jumped from humans to mice, rapidly accumulated mutations conducive to infecting that host, then jumped back into humans, indicating an inter-species evolutionary trajectory for the Omicron outbreak.

## INTRODUCTION

The coronavirus disease 2019 (COVID-19) pandemic, caused by the SARS-CoV-2 RNA virus, has led to significant illness and death worldwide. The SARS-CoV-2 Omicron variant was first reported in South Africa on November 24^th^, 2021, and was designated as a variant of concern (VOC) within two days by the World Health Organization (WHO) based on the increase in infections attributable to this variant in South Africa (*i*.*e*., Omicron outbreak). In addition, the open reading frame encoding the spike protein (ORF *S*) of Omicron harbors an exceptionally high number of mutations. These mutations are particularly relevant to SARS-CoV-2 infection characteristics because the spike protein is well-known to mediate viral entry into the host cell by interacting with angiotensin-converting enzyme 2 (ACE2) on the cell surface (Zhou et al., 2020). In addition, the spike protein is also a target for vaccine development and antibody-blocking therapy (Huang et al., 2020; Martinez-Flores et al., 2021).

The proximal origins of Omicron have quickly become a controversial topic of heated debate in the scientific and public health communities (Callaway, 2021; Kupferschmidt, 2021). Many mutations detected in Omicron were rarely reported among previously sequenced SARS-CoV-2 variants (Shu and McCauley, 2017; Hadfield et al., 2018), leading to three prevalent hypotheses regarding its evolutionary history. The first hypothesis is that Omicron could have “cryptically spread” and circulated in a population with insufficient viral surveillance and sequencing. Second, Omicron could have evolved in a chronically infected COVID-19 patient, such as an immunocompromised individual who provided a suitable host environment conducive to long-term virus adaptation. The third possibility is that Omicron could have accumulated mutations in a nonhuman host and then jumped into humans. Currently, the second scenario represents the most popular hypothesis regarding the proximal origins of Omicron (Callaway, 2021; Kupferschmidt, 2021).

The first two hypotheses assume that Omicron acquired these mutations in humans (collectively to as “human origin hypothesis” hereafter), while the third assumes that Omicron acquired mutations in a nonhuman species. Based on our previous work in viral evolution, we hypothesized that the host species in which Omicron acquired its particular set of mutations could be determined by analyzing the molecular spectra of mutations (*i*.*e*., the relative frequency of the twelve types of base substitutions). In previous work, we showed that *de novo* mutations in RNA virus genomes are generated in a replication-independent manner and are highly dependent on mutagenic mechanisms specific to the host cellular environment, resulting in overrepresentation with specific mutation types. For example, reactive oxygen species (ROS) can oxidize guanine to 8-oxoguanine and thereby induce the G>U transversion (Li et al., 2006; Kong and Lin, 2010), while cytidine deaminases can induce RNA editing such as C>U transitions (Blanc and Davidson, 2010; Harris and Dudley, 2015). Consistent with this phenomenon, viruses belonging to different orders (*e*.*g*., poliovirus, Ebola virus, and SARS-CoV-2) were found to exhibit similar molecular spectra of mutations when evolving in the same host species, while members of the same virus species exhibit divergent molecular spectra when evolving in different host species (Shan et al., 2021). Since *de novo* mutations can thus strongly influence the molecular spectrum of mutations that accumulate during virus evolution in a host-specific manner, the host species in which Omicron acquired its mutations could be determined by analyzing information carried by the mutations themselves.

In this study, we identified mutations acquired by Omicron before its outbreak, and tested whether the molecular spectrum of these mutations was consistent with the cellular environment of human hosts. Prominent dissimilarities were observed between the molecular spectrum of Omicron and a relatively comprehensive set of molecular spectra from variants known to have evolved in humans, including those of three isolates from chronic SARS-CoV-2 patients. Therefore, we next examined the molecular spectra of mutations obtained from a wide range of host mammals for comparison with that of Omicron. Finally, we used molecular docking-based analyses to investigate whether the mutations in the Omicron spike protein could be associated with adaptation to the host species inferred from molecular spectrum analysis. Our study provides insight into the evolutionary trajectory and proximal origins of Omicron through careful scrutiny of its mutations, and suggests strategies for avoiding future outbreaks caused by potentially dangerous SARS-CoV-2 variants.

## RESULTS

### Over-representation of nonsynonymous mutations in Omicron ORF *S* suggests strong positive selection

To first identify mutations that accumulated in the SARS-CoV-2 genome prior to the Omicron outbreak, we constructed a phylogenetic tree that included the genomic sequences of the reference SARS-CoV-2, two variants in the B.1.1 lineage which were genetically close to Omicron, and 48 Omicron variants sampled before November 15^th^, 2021 (**Fig. 1A**). These two B.1.1 variants were sampled during April 22^nd^–May 5^th^, 2020, which suggested that the progenitor of Omicron diverged from the B.1.1 lineage roughly in mid-2020. Intermediate versions have gone largely undetected, thus resulting in an exceptionally long branch leading to the most recent common ancestor (MRCA) of Omicron in the phylogenetic tree (**Fig. 1A**). We hereafter refer to this long branch as Branch O.

**Fig. 1.**
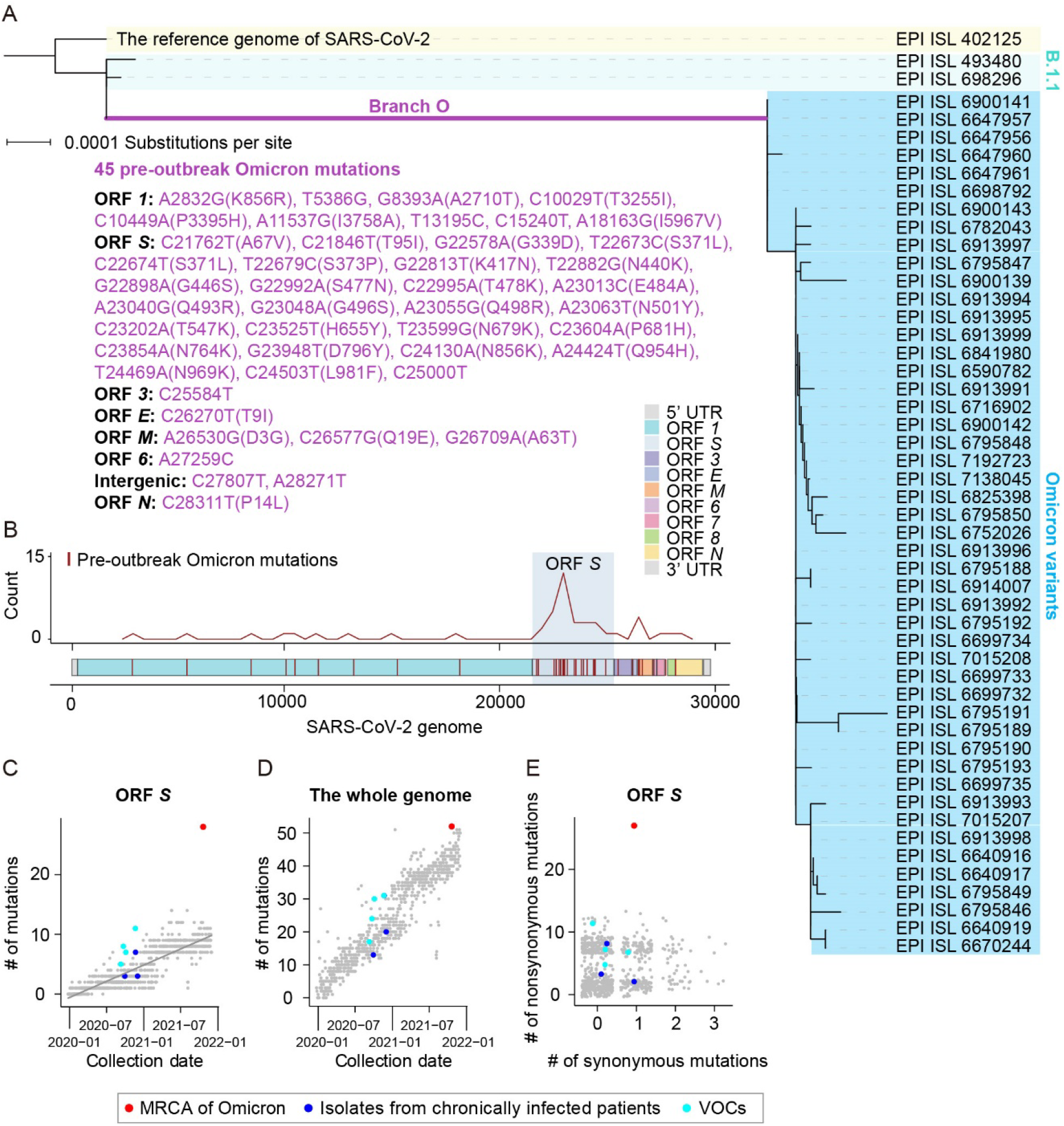
The characterization of pre-outbreak Omicron mutations. **A**. The phylogenetic tree of Omicron variants, including the reference genome of SARS-CoV-2 (EPI_ISL_402125), two B.1.1 variants, and 48 Omicron variants. A total of 45 pre-outbreak Omicron point mutations in the long branch leading to the MRCA of Omicron (Branch O, labeled in purple) in the phylogenetic tree are grouped according to ORFs. **B**. The distribution of pre-outbreak Omicron mutations across the SARS-CoV-2 genome. The curve indicates the density of mutations. UTR stands for the untranslated region. **C**. Number of mutations that accumulated in ORF *S* of the MRCA of current Omicron variants (red), the other four VOCs (*i*.*e*., Alpha, Beta, Gamma, and Delta; cyan), and three SARS-CoV-2 isolates from chronically infected patients (blue), against the date of sample collection. SARS-CoV-2 variants randomly sampled (one variant per day) are shown in grey, and the grey line represents their linear regression. **D**. Similar to (C), for the whole genome. **E**. A scatterplot shows the numbers of synonymous and nonsynonymous mutations in ORF *S* (jittered in order to reduce overplotting).

We identified 45 point mutations that were introduced in Branch O (hereafter referred to as “pre-outbreak Omicron mutations”; **Fig. 1A**). Visual assessment suggested that the pre-outbreak Omicron mutations were over-represented in ORF *S* (**Fig. 1B**). To test if the rate at which mutations accumulated in ORF *S* was accelerated in Branch O, we randomly sampled one SARS-CoV-2 variant per day since December 24^th^, 2019 from the Global Initiative on Sharing All Influenza Data (GISAID) (Shu and McCauley, 2017) to compare mutation accumulation rates among different variants. We found that mutations accumulated in ORF *S* at a rate of ∼0.45 mutations per month on average. In sharp contrast, 27 mutations accumulated in ORF *S* in Branch O during the 18 months spanning May 2020–November 2021, equivalent to ∼1.5 mutations per month, or ∼3.3 times faster than the average rate of other variants (**Fig. 1C**).

Counting mutations across the whole SARS-CoV-2 genome indicated that Omicron acquired mutations in the genome at a similar rate to other variants (**Fig. 1D**), suggesting that the accelerated evolutionary rate of ORF *S* could not be explained by an overall elevated mutation rate in Omicron progenitors. In light of these findings, we hypothesized that positive selection could have helped accelerate the evolutionary rate of ORF *S*. To test this hypothesis, we sought to infer the strength of positive selection by estimating the ratio of nonsynonymous to synonymous mutations. Twenty-six of the 27 pre-outbreak mutations in the ORF *S* of Omicron were nonsynonymous (**Fig. 1E**), resulting in a *d*_N_/*d*_S_ ratio of 6.64, significantly greater than a *d*_N_/*d*_S_ of 1.00 (*P* = 0.03, Fisher’s exact test). These results indicated that positive selection contributed to increasing the mutation rate in ORF *S* in Branch O.

To test if such a level of positive selection is common among SARS-CoV-2 variants, we counted the number of nonsynonymous and synonymous mutations in ORF *S* in other SARS-CoV-2 VOC lineages (*i*.*e*., Alpha, Beta, Gamma, and Delta) as well as in the genomes of SARS-CoV-2 variants isolated from three chronically infected patients (Kemp et al., 2021; Truong et al., 2021). None of these other VOCs or isolates exhibited comparable numbers of nonsynonymous mutations as that of mutations in Branch O (**Fig. 1E**). These observations strongly suggested that the Omicron variant had undergone a strong positive selection for the spike protein that no other known SARS-CoV-2 variants evolved in humans had been subjected to. Considering that the spike protein determines the host range of a coronavirus (*i*.*e*., which organisms it can infect), we therefore hypothesized that the progenitor of Omicron might host-jump from humans to a nonhuman species because this process would require substantial mutations in the spike protein for rapid adaptation to a new host.

### The molecular spectrum of pre-outbreak Omicron mutations is inconsistent with an evolutionary history in humans

Previous studies showed that the molecular spectrum of mutations that accumulate in a viral genome reflect a host-specific cellular environment (Deng et al., 2021; Shan et al., 2021). To test the human origin hypothesis of Omicron, we compared the molecular spectrum of the 45 pre-outbreak Omicron mutations with the “standard” molecular spectrum for SARS-CoV-2 variants known to have evolved strictly in humans (hereafter referred to as “the hSCV2 spectrum”, **Fig. 2A**). The hSCV2 spectrum included 6,986 point mutations that were compiled from 34,853 high-quality sequences of SARS-CoV-2 variants isolated from patients worldwide (Shan et al., 2021). We found that the molecular spectrum of the pre-outbreak Omicron mutations was significantly different from the hSCV2 spectrum (*P* = 0.004, *G*-test, **Fig. 2B**). In particular, transitions were more abundant than transversions and C>U mutation was more abundant than its complementary mutation G>A, as in the hSCV2 spectrum; However, a hallmark of RNA virus mutations when evolving in humans—a higher rate of G>U mutation than its complementary mutation C>A (Panchin and Panchin, 2020; De Maio et al., 2021; Deng et al., 2021; Shan et al., 2021), likely caused by cellular ROS—was absent in the pre-outbreak Omicron mutations.

**Fig. 2.**
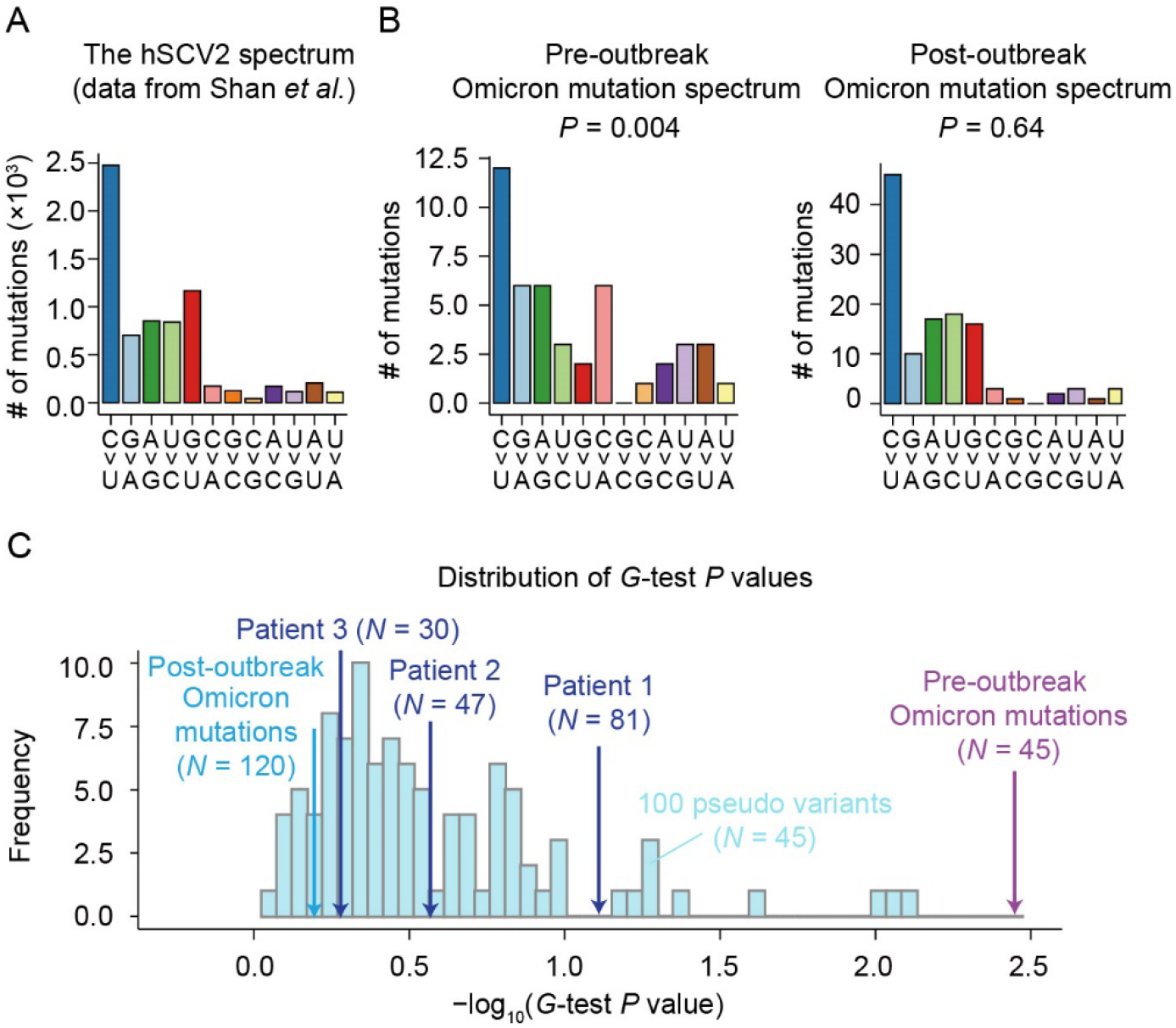
Comparison of the molecular spectrum of pre-outbreak Omicron mutations and spectra of mutations known to accumulate in humans. **A**. The molecular spectrum of viral mutations that accumulated in humans (the hSCV2 spectrum). **B**. The molecular spectra of pre-and post-outbreak Omicron mutations. *P* values were given by *G-*test to test whether a molecular spectrum was significantly different from the hSCV2 spectrum. **C**. The distribution of *P* values (given by *G*-test) of 100 pseudo samples that were down sampled from the hSCV2 spectrum. The number of mutations (*N*) of each pseudo sample was equal to 45. The molecular spectra of SARS-CoV-2 isolates from three chronically infected patients were also labeled. SARS-CoV-2 data of patient 1 were retrieved from Kemp *et al*. (2021) and those of patients 2 and 3 were retrieved from Truong *et al*. (2021).

To exclude the possibility that this apparent difference in the molecular spectrum was caused by the relatively small number of pre-outbreak Omicron mutations, we generated 100 “pseudo” variants *in silico* by randomly down sampling 45 mutations from the hSCV2 spectrum. None of the pseudo variants showed smaller *P* values (based on *G*-tests) than that obtained using the pre-outbreak Omicron mutations (**Fig. 2C**), nor did the SARS-CoV-2 isolates with mutations known to be acquired in the three chronically infected patients (# of mutations are 30, 47, and 81, **Fig. 2C**). These observations indicated that the difference between the molecular spectrum of pre-outbreak Omicron mutations and the hSCV2 spectrum could not be strictly attributed to statistical randomness.

To exclude the possibility that some mutations which occurred early in the evolution of Omicron distorted the molecular spectrum of mutations that accumulated afterward, we identified 120 point mutations on top of the MRCA of Omicron, by screening 695 Omicron variants collected spanning November 8^th^–December 7^th^, 2021 (hereafter referred to as “post-outbreak Omicron mutations”). The molecular spectrum of these post-outbreak Omicron mutations was not significantly different from the hSCV2 spectrum (**Fig. 2B–C**). This finding indicated that Omicron acquired mutations following the same molecular spectrum as other SARS-CoV-2 variants during its evolution in human hosts. Collectively, these molecular spectrum analyses revealed that pre-outbreak Omicron mutations were unlikely to have been acquired in humans.

### The molecular spectrum of pre-outbreak Omicron mutations is consistent with an evolutionary history in mice

In light of our findings that Omicron may have evolved in another host before its outbreak, we next sought to determine the nonhuman host species in which these mutations accumulated. To this end, we first characterized the molecular spectra of coronaviruses that evolved in different host species for comparison with that of Omicron. Specifically, we retrieved 17 sequences of murine hepatitis viruses, 13 canine coronaviruses, 54 feline coronaviruses, 23 bovine coronaviruses, and 110 porcine deltacoronaviruses (**Table S1**), constructed the phylogenetic tree for the coronaviruses isolated from each host species (canine coronavirus as an example shown in **Fig. 3A** and the rest are shown in **Fig. S1**), and identified the mutations that accumulated in each branch (**Fig. 3A**). We also included some previously reported molecular spectra (Shan et al., 2021), including 17 spectra of mutations acquired by SARS-CoV-, SARS-CoV-2-, and MERS-CoV-related coronaviruses during their evolution in bats, two spectra of camel MERS-CoV, one spectrum estimated from 807 MERS-CoV mutations accumulated in human (the hMERS spectrum), as well as the hSCV2 spectrum. Furthermore, we also included the molecular spectrum of mutations identified in an early variant of each of the other four VOCs.

**Fig. 3.**
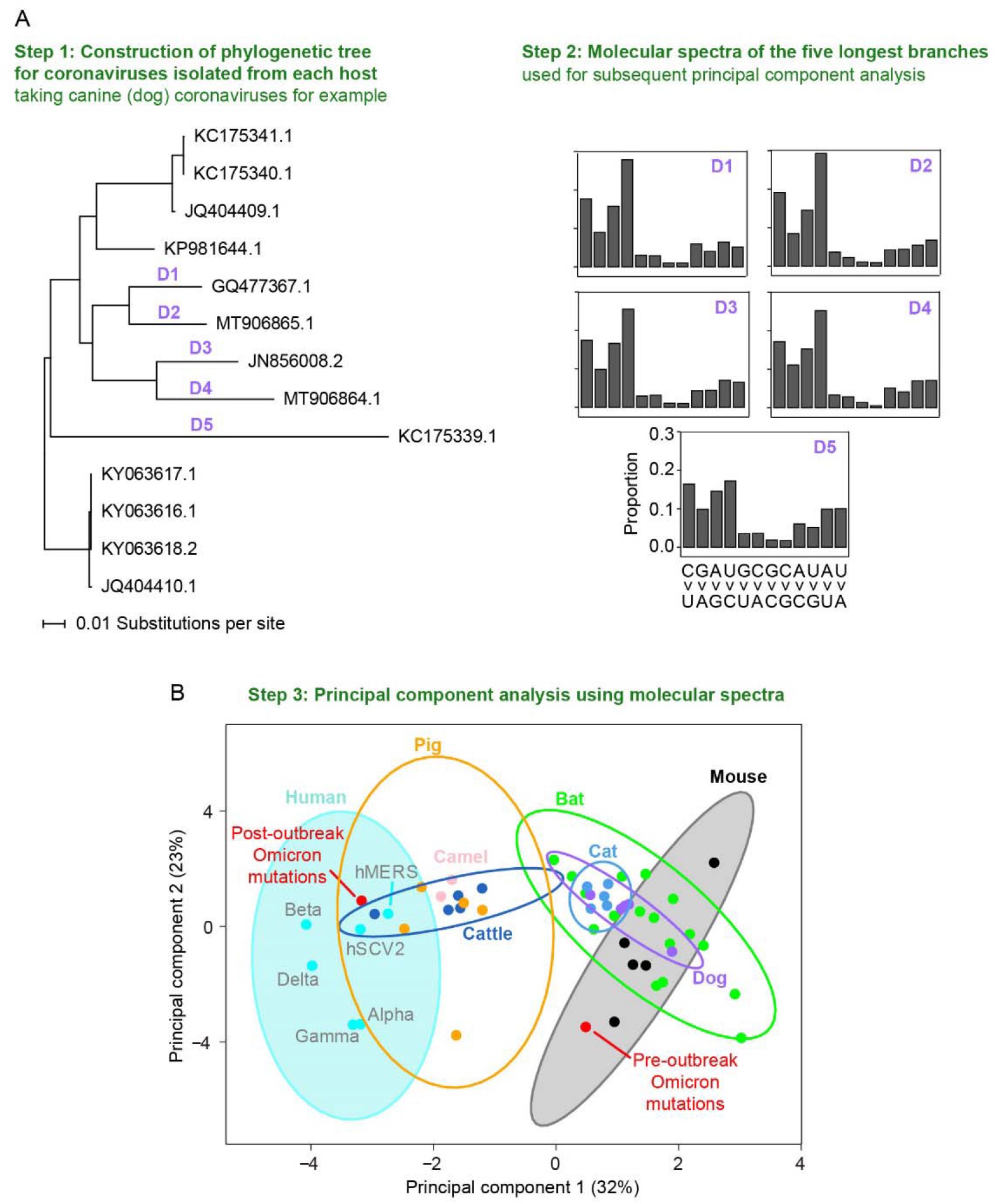
The similarity in molecular spectra between Omicron and coronaviruses isolated from various mammalian species. **A**. A schematic shows the workflow for analyzing the similarity in molecular spectra across various hosts, taking variants in dogs as an example. **B**. The principal component analysis plot depicts the molecular spectra of virus mutations that accumulated in humans and various host species. Dots were colored according to the corresponding host species. The 95% confidence ellipses are shown for each host species.

We performed principal component analysis to reduce the dimensionality of the molecular spectrum of mutations, and subsequently visualized the data using the first two principal components (**Fig. 3B**). Consistent with the results of our previous study (Shan et al., 2021), drawing 95% confidence ellipses for each host species showed that the molecular spectra clustered according to their respective hosts (**Fig. 3B**), likely because viruses evolving in the same host species share the mutagens specific to that host’s cellular environment. Supporting this point, the molecular spectrum of post-outbreak Omicron mutations (which are known to have accumulated in humans) was located within the human 95% confidence ellipse. In contrast, the molecular spectrum of pre-outbreak Omicron mutations was within the mouse ellipse, suggesting that the pre-outbreak mutations accumulated in a rodent (in particular mouse) host.

### Pre-outbreak Omicron mutations in the spike protein significantly overlap with mutations in mouse-adapted SARS-CoV-2

Mice were previously reported to serve as poor hosts for SARS-CoV-2 because the spike protein of early SARS-CoV-2 variants exhibit low-affinity interactions with mouse ACE2 (Lam et al., 2020; Zhou et al., 2020; Ren et al., 2021; Wong et al., 2021). However, over the course of the pandemic SARS-CoV-2 variants emerged that could infect mice. For example, variants harboring the spike mutation N501Y, which are relatively common in human patients (24.7%), could infect mice (Gu et al., 2020; Leist et al., 2020; Sun et al., 2021). If the progenitor of Omicron indeed evolved in a mouse species before the Omicron outbreak, we postulated that its spike protein likely adapted through increased binding affinity for mouse ACE2. To test this possibility, we projected the pre-outbreak Omicron mutations in the spike protein onto a three-dimensional structural model of the spike:ACE2 complex (Lan et al., 2020). Seven mutations (*i*.*e*., K417N, G446S, E484A, Q493R, G496S, Q498R, and N501Y) were located at the interface of ACE2 and the receptor-binding domain (RBD) of the spike protein and could potentially affect their interactions (**Fig. 4A**).

**Fig. 4.**
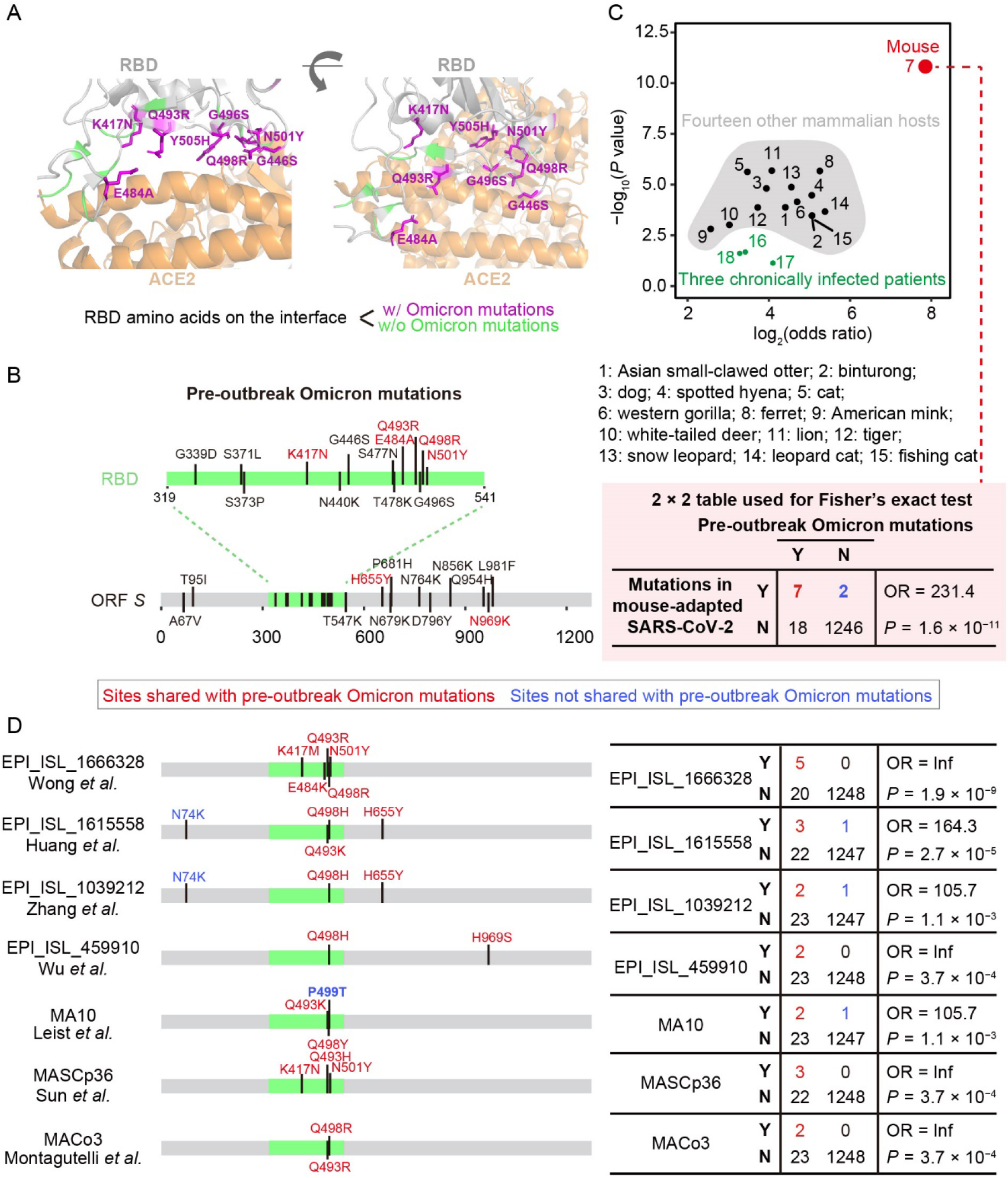
The similarity in the spike protein sequence between Omicron and SARS-CoV-2 variants isolated from various hosts. **A**. The structure of the interface between SARS-CoV-2 spike protein and human ACE2, from the crystal structure of the spike:ACE2 (human) complex (PDB: 6M0J). RBD residues on the interface (defined within 5□ distance) were colored. **B**. The amino acid mutations in the spike protein in the MRCA of Omicron variants. **C**. The statistical assessment on the overlapping in mutated positions between Omicron and SARS-CoV-2 variants using Fisher’s exact test. The 2 × 2 contingency table for mice is shown. OR stands for the odds ratio. **D**. Comparison between pre-outbreak Omicron mutations and mutations detected in seven SARS-COV-2 variants isolated from mice, in the spike protein. RBD of spike protein was colored green.

Previous studies of SARS-CoV-2 variants isolated from mice reported specific amino acid mutations in the spike protein that could promote its interactions with mouse ACE2 (Leist et al., 2020; Wu et al., 2020b; Huang et al., 2021; Montagutelli et al., 2021; Sun et al., 2021; Wong et al., 2021; Zhang et al., 2021). In addition, previous studies have described some reverse zoonotic events (*e*.*g*., from humans to other mammals such as mink and white-tailed deer) for SARS-CoV-2 (Chandler et al., 2021; Oude Munnink et al., 2021), and the variants isolated from these mammalian hosts presumably harbored amino acid mutations that could potentially participate in their adaptation to these hosts. Thus, if the progenitor of Omicron evolved in mice and adapted to mouse ACE2, we predicted that the pre-outbreak Omicron mutations should share considerable overlap with the mutations identified in these mouse-adapted SARS-CoV-2 variants, but not those of other mammalian species.

To test this prediction, we identified the mutations in ORF *S* of SARS-CoV-2 variants isolated from 15 mammalian species (*e*.*g*., mice, cats, dogs, minks, and deer, **Table S2**) and found that pre-outbreak Omicron mutations tended to share the same positions as the ORF *S* mutations identified in mice (odds ratio = 231.4, *P* = 1.6 × 10^−11^, Fisher’s exact test, **Fig. 4B–C**). In contrast, same statistical test showed much lower odds ratios and significance levels for overlap in these mutations with other species (**Fig. 4C)**. Pre-outbreak Omicron mutations also overlapped with some mutations detected in isolates from chronically infected patients (Kemp et al., 2021; Truong et al., 2021), although they too showed substantially lower odds ratios and significance levels (**Fig. 4C**). These observations implied that the pre-outbreak Omicron mutations in ORF *S* promoted its adaptation to a mouse host.

We then conducted enrichment analysis for each of the seven mouse-adapted SARS-CoV-2 variants and observed statistical significance for all these variants (**Fig. 4D**). In particular, we observed amino acid mutations at residues 493 and 498 in five and six of the seven mouse-adapted SARS-CoV-2 variants, respectively (**Fig. 4D**). Identical amino acid mutations (*i*.*e*., Q493R and Q498R) were both observed in two variants (Montagutelli et al., 2021; Wong et al., 2021), and considering that these two amino acid mutations are uncommon in human patients infected by non-Omicron SARS-CoV-2 variants (0.005% and 0.002%, respectively) we concluded that the progenitor of Omicron evolved in mouse species (or at least rodent species).

### Pre-outbreak Omicron mutations in the spike protein significantly enhance binding affinity with mouse ACE2

To investigate the mechanisms by which the pre-outbreak Omicron mutations in the spike protein could have contributed to its adaptation to a mouse host, we examined their interaction through molecular docking analysis of the spike protein RBD and mouse ACE2 (**Fig. 5A**). Following previous studies (Lam et al., 2020; Rodrigues et al., 2020), we estimated the HADDOCK score (van Zundert et al., 2016), which is positively associated with the dissociation constant (*K*_D_, with smaller *K*_D_ indicating stronger binding) of protein interactions (Kastritis and Bonvin, 2010), and can be used to predict the susceptibility of a mammalian species to infection with SARS-CoV-2 (Rodrigues et al., 2020).

**Fig. 5.**
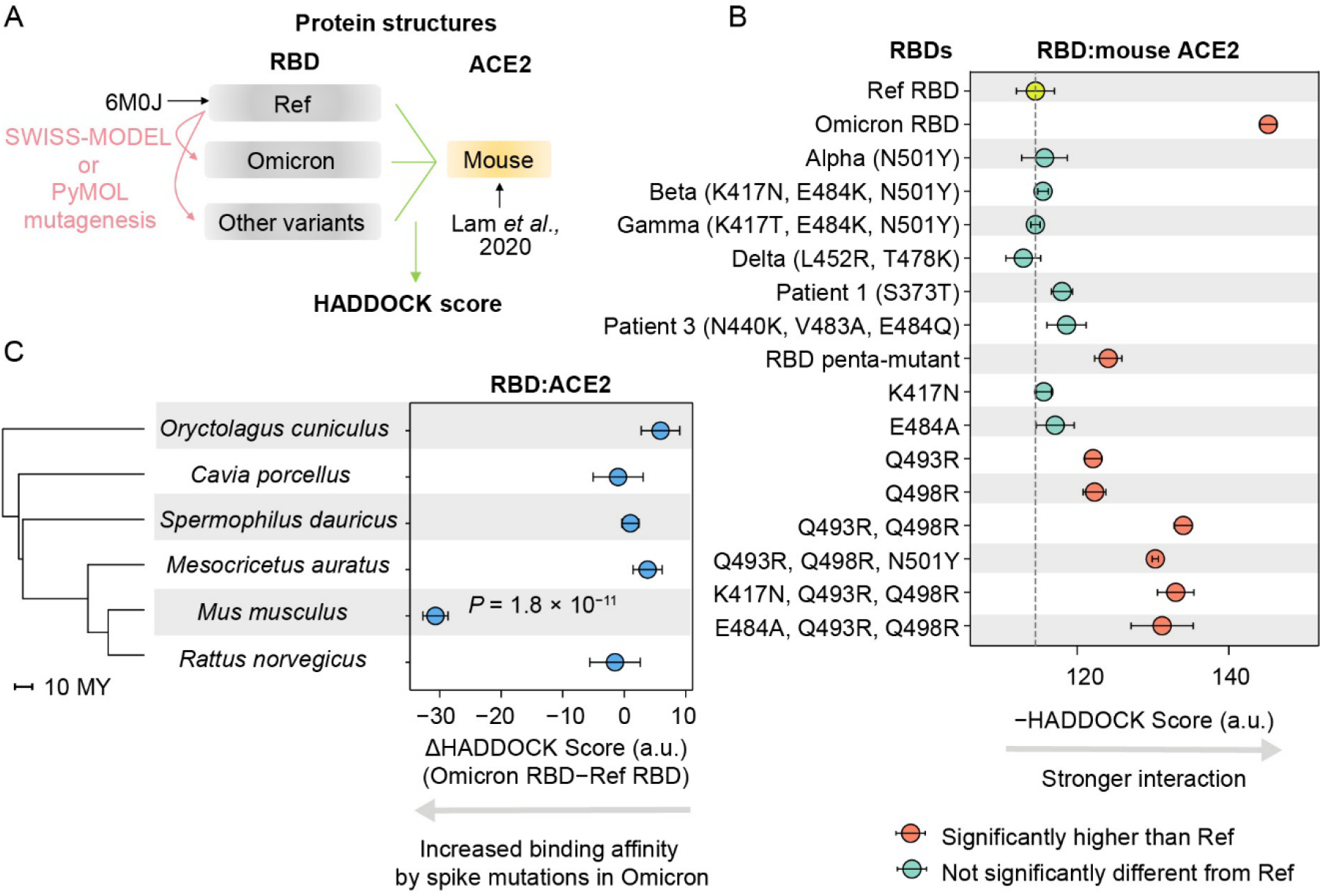
Predicted binding affinities between RBD variants and ACE2. **A**. A schematic shows the workflow to estimate the HADDOCK scores between RBD variants and mouse ACE2. **B**. The HADDOCK scores for various RBD variants and mouse ACE2. The error bars represent standard errors. Penta-mutant of RBD harbored five mutations (K417N, E484A, Q493R, Q498R, and N501Y). Patient 2 who did not harbor any amino acid mutations in RBD was not shown. **C**. The ΔHADDOCK scores for five rodent species and the European rabbit. *P* values were given by two-tailed *t*-tests. Only *P* values <0.05 are labeled. The phylogenetic tree was constructed using TIMETREE, in the unit of million years (MY).

To confirm the accuracy of inferences regarding the binding affinity between spike protein RBD and ACE2 based on the HADDOCK score, we calculated the HADDOCK score for eight experimentally determined *K*_D_ values between four RBD variants and human (or mouse) ACE2 (Sun et al., 2021). The HADDOCK scores were positively correlated with the *K*_D_ values in the analysis (Pearson’s correlation coefficient *r* = 0.93, *P* = 0.002, **Fig. S2A–C**), thus supporting the validity of molecular docking-based predictions of ACE2-binding affinity for other RBD variants.

The molecular docking-based predictions suggested that the RBD of Omicron exhibited higher binding affinity for mouse ACE2 than that of RBD encoded in the reference SARS-CoV-2 genome, further suggesting an evolutionary history in mice (**Fig. 5B**). And as expected, the mutations detected in the RBD of the other four VOCs of SARS-CoV-2 as well as those of variants isolated from chronically infected human patients showed no apparent changes in their binding affinity for mouse ACE2 compared with the reference RBD (**Fig. 5B**).

Since five amino acid mutations were shared between Omicron and mouse-adapted SARS-CoV-2 variants in RBD (*i*.*e*., K417N, E484A, Q493R, Q498R, and N501Y; **Fig. 4B**), and that they together enhanced RBD binding affinity for mouse ACE2 (**Fig. 5B**), we next determined the individual effects of each of these five mutations. Notably, only Q493R and Q498R significantly increased the binding affinity with mouse ACE2, which was consistent with their repeated detection in mouse-adapted SARS-CoV-2 variants (Montagutelli et al., 2021; Wong et al., 2021). Indeed, docking analysis showed that Q493R/Q498R double mutation could further increase the RBD binding affinity for mouse ACE2 (**Fig. 5B**). By contrast, the other three mutations showed no significant effects on the binding affinity between RBD and mouse ACE2, neither in the reference RBD nor in the Q493R/Q498R double mutant (**Fig. 5B**), suggesting that they did not contribute to the enhanced interaction between Omicron RBD and mouse ACE2. Indeed, previous studies showed that K417N, E484K, and N501Y were related to escape from neutralizing antibodies (Li et al., 2021; Nelson et al., 2021).

### Mice as the most likely rodent species in which the progenitor of Omicron evolved

While the observations regarding both the molecular spectrum of mutations and the RBD-ACE2 interaction suggested that mice were candidate host species in which the progenitor of Omicron evolved, it remained plausible that Omicron evolved in some other rodent species with similar cellular mutagen environment and ACE2 structure to mice. We postulated that if Omicron evolved in another rodent species, the amino acid mutations in Omicron RBD should elevate its interaction with the ACE2 of this host. To test this prediction, we applied molecular docking analysis to four additional rodent species representing different lineages of rodents (Kumar et al., 2017)—brown rats (*Rattus norvegicus*), guinea pigs (*Cavia porcellus*), golden hamsters (*Mesocricetus auratus*), and Daurian ground squirrels (*Spermophilus dauricus*)—as well as a close relative of rodents, European rabbits (*Oryctolagus cuniculus*). Omicron RBD showed higher ACE2-binding affinity (compared with the reference RBD) only to the mouse ACE2 (**Fig. 5C**), suggesting that mice are the most likely host species in which the progenitor of Omicron evolved.

## DISCUSSION

In this study, we used the molecular spectrum of mutations of the SARS-CoV-2 Omicron variant to trace its proximal host origins. We found that the molecular spectrum of pre-outbreak Omicron mutations was inconsistent with the rapid accumulation of mutations in humans, but rather suggested a trajectory in which the progenitor of Omicron experienced a reverse zoonotic event from humans to mice sometime during the pandemic (most likely in mid-2020) and accumulated mutations in a rodent host (most likely mouse) for more than one year before jumping back to humans in late-2021. While evolving in mice, the progenitor of Omicron adapted to the mouse host by acquiring amino acid mutations in the spike protein that increased its binding affinity with mouse ACE2. In addition, mutations associated with immune escape also accumulated, which may also be a contributing factor in its rapid spread.

While we show a phylogenetically long branch leading to the MRCA of current Omicron variants (*i*.*e*., Branch O), it is worth noting that intermediate versions of Omicron were occasionally reported. For example, a SARS-CoV-2 variant (EPI_ISL_7136300) was collected by the Utah Public Health Laboratory on December 1^st^, 2021 which harbored 32 of the 45 pre-outbreak Omicron mutations. However, the 13 mutations absent in this variant clustered within residues 371–501 of the spike protein (**Fig. S3**). The absence of these spike protein mutations thus suggested that this variant was a product of recombination between an Omicron variant and another SARS-CoV-2 variant, rather than a direct progenitor of Omicron. Considering the large number of pre-outbreak Omicron mutations (45) combined with the sparsity of intermediate versions identified to date, this long branch leading to Omicron in our phylogenetic reconstruction remains valid.

Although we primarily focused on point mutations because the molecular spectrum of these mutations can reflect the host cellular environment (Deng et al., 2021; Shan et al., 2021), we also used the information of deletions and insertions to infer the evolutionary trajectory of Omicron. For example, a B.1.1 variant (EPI_ISL_493480) shared the same deletion (Δ105–107 in non-structural protein 6) as the Omicron variants, which was used to infer that B.1.1 is a close relative of Omicron. In addition, spike Δ69–70 deletion is shared by Omicron and many non-Omicron variants isolated from patients (Meng et al., 2021), but is absent in the early samples of SARS-CoV-2 (Wu et al., 2020a), strongly suggesting that the progenitor of Omicron was jumped from humans to mice during the pandemic.

In addition, we noted that Omicron harbored a nine nucleotide insertion (GAGCCAGAA, encoding the peptide EPE) after residue 214 in the spike protein. This insertion is identical to the sequence of *TMEM245* in the human genome or that of ORF *S* in the human coronavirus hCoV-229E, which was used as evidence to support a human origin for Omicron (Venkatakrishnan et al., 2021). However, we provide a simpler explanation for this insertion, namely that it was derived from an RNA fragment of ORF *N* in the SARS-CoV-2 genome (**Fig. S4**). We believe that the insertion of an ORF *N* fragment is more likely because the RNA abundance of ORF *N* is much higher than that of mRNA encoded by the human genome (Wei et al., 2021). That is, in SARS-CoV-2-infected cells, a substantial proportion of RNAs are viral, and especially so for ORF *N* due to the nested nature of the coronavirus genome and subgenomes (Kim et al., 2020).

It also warrants mention that all of the mouse-adapted SARS-CoV-2 variants were amplified/purified in Vero cells (a cell line originally isolated from the kidney of green monkey) at some stage of experimentation, which could impose an additional selection pressure to enhance the spike protein binding affinity towards primate ACE2 (Leist et al., 2020; Wu et al., 2020b; Huang et al., 2021; Montagutelli et al., 2021; Sun et al., 2021; Wong et al., 2021; Zhang et al., 2021). Consistent with this experimental process, the amino acid mutations acquired by Omicron and mouse-adapted viruses were not always identical, even if mutations occurred at the same residue. For example, Q493H and Q493K were also detected in the mouse-adapted SARS-CoV-2 at residue 493 of the spike protein, in addition to mutations observed in Omicron (Q493R). Different from the effects of Q493R, these two mutations increased the binding affinity toward both mouse and human ACE2 (**Fig. S2C**), indicating that SARS-CoV-2 could potentially evolve remarkably high diversity in its adaptation to ACE2 from various host species. Consistent with this possibility, numerous mutations were also identified in the spike protein of SARS-CoV-2 RNA fragment amplified from wastewater samples (Smyth et al., 2021).

Humans represent the largest known reservoir of SARS-CoV-2. Our study suggests that SARS-CoV-2 could have spilled over from humans to wild animals, and that the variants which successfully infected animal hosts could then accumulate new mutations before jumping back into humans as a variant of concern. Given the ability of SARS-CoV-2 to jump across various species, it appears likely that global populations will face additional animal-derived variants until the pandemic is well under control. Viral surveillance and sequencing in wild animals will likely help to prevent future outbreaks of dangerous SARS-CoV-2 variants.

## METHODS

### Identification of pre-outbreak and post-outbreak Omicron mutations

Genomic sequences of 695 SARS-CoV-2 Omicron variants were downloaded from GISAID (https://www.gisaid.org/) on December 7th, 2021. The reference genome of SARS-CoV-2 (EPI_ISL_402125) and two variants in the B.1.1 lineage (EPI_ISL_698296 and EPI_ISL_493480) were also downloaded from GISAID.

The genomes of SARS-CoV-2 variants were aligned by MUSCLE v3.8.1551 (Edgar, 2004). The phylogenetic tree and ancestral sequences were reconstructed using FastML v3.11 (Ashkenazy et al., 2012) with default parameters. The single-nucleotide substitutions obtained by the most recent common ancestor (MRCA) of Omicron variants after its divergence from the B.1.1 lineage were defined as pre-outbreak Omicron mutations. To detect the post-outbreak Omicron mutations, the sequences of 695 Omicron variants were aligned to the Omicron’s MRCA sequence, and sequences with >10 single-nucleotide substitutions were discarded. The single-nucleotide substitutions detected in at least two variants were defined as the post-outbreak Omicron mutations.

The numbers of synonymous and nonsynonymous sites in ORF *S* of SARS-CoV-2 were estimated by PAML in a previous study (Wei et al., 2021). Briefly, *d*_N_ was calculated as the ratio between the number of nonsynonymous mutations and the number of nonsynonymous sites, while *d*_S_ was calculated as the ratio between the number of synonymous mutations and the number of synonymous sites.

The frequencies of three mutations (Q493R, Q498R, and N501Y) among patients were retrieved from CoV-GLUE-Viz (http://cov-glue-viz.cvr.gla.ac.uk/) updated at November 23^th^, 2021.

### Comparison between the sequence evolutionary rate of Omicron and other SARS-CoV-2 variants

A total of 764 variant sequences were randomly sampled from the SARS-CoV-2 genomic sequences deposited at GISAID, one variant each day since COVID-19 outbreak. The progenitors of other four VOCs (Alpha, Beta, Gamma, and Delta) were retrieved from Nextstrain (https://nextstrain.org/) (Hadfield et al., 2018). Single-nucleotide substitutions (relative to the reference genome) of each variant were defined as the mutations acquired by the SARS-CoV-2 variant. The single-nucleotide base substitutions of three chronically infected patients were retrieved from two previous studies (Kemp et al., 2021; Truong et al., 2021). The mutations with allele frequency >50% on the final monitored day were used to count mutations that accumulated in a chronically infected patient.

We performed resampling test to estimate the statistical significance. Specifically, we randomly sampled 45 mutations from the 6,986 point mutations identified in a previous study from the 34,853 high-quality sequences of SARS-CoV-2 variants isolated from patients worldwide (Shan et al., 2021). This operation was repeated 100 times *in silico*.

### Characterization of molecular spectra of mutations

Complete genomic sequences of 23 bovine coronavirus (*Betacoronavirus 1*), 13 canine coronavirus (*Alphacoronavirus 1*), 54 feline coronavirus (*Alphacoronavirus 1*), 17 murine hepatitis virus (*Murine coronavirus*), and 110 porcine deltacoronavirus (*Coronavirus HKU15*) were downloaded from National Center for Biotechnology Information (NCBI) Virus database (https://www.ncbi.nlm.nih.gov/labs/virus/vssi/) (Hatcher et al., 2017), querying the hosts as *Bos taurus* (cattle), *Canis lupus familiaris* (dogs), *Felis catus* (cats), *Mus musculus* (mice), and *Sus scrofa* (pigs), respectively (**Table S1**).

The virus genome sequences were aligned by MUSCLE, and the phylogenetic trees and ancestral sequences were reconstructed using FastML. Since the roots of these phylogenetic trees were not readily identified, we kept only external branches to ensure the correction direction of base substitutions (*e*.*g*., C>U vs. U>C). For the sake of clarity, we showed the molecular spectra for five branches with the largest number of mutations for each coronavirus species in the main text. The full data set is available in **Table S1**.

We characterized the molecular spectra of mutations accumulated in chronically infected patients, in which single-nucleotide base substitutions that ever occurred during the monitored period were counted. We downloaded the genomic sequences of four variants (EPI_ISL_5803018, EPI_ISL_3730369, EPI_ISL_4003132, and EPI_ISL_6260720), each from one of the other four VOCs (Alpha, Beta, Gamma, and Delta, respectively), to estimate the molecular spectra of mutations accumulated in VOCs.

### Principal component analyses

We performed principal component analysis (prcomp function in *R*) with the proportions of the 12 base-substitution types as the input, and then projected molecular spectra into a two-dimensional space according to the first two principal components. To define the borderlines of molecular spectra for each host species (*i*.*e*., cattle, bats, dogs, cats, mice, pigs, or humans), we estimated the 95% confidence ellipses (stat_ellipse option in *R*) from the molecular spectra of these host species. The spectra of pre-and post-outbreak Omicron mutations were further projected into the same two-dimensional space.

### Comparison of pre-outbreak Omicron mutations with mutations detected in SARS-COV-2 variants isolated from various mammalian hosts

We downloaded from GISIAD the genomic sequences of SARS-CoV-2 variants isolated from 18 mammalian hosts (**Table S2**): *Aonyx cinereus* (Asian small-clawed otter); *Arctictis binturong* (binturong); *Canis lupus familiaris* (dog); *Crocuta crocuta* (spotted hyena); *Felis catus* (cat); *Gorilla gorilla* (western gorilla); *Mus musculus* (mouse); *Mustela furo* (ferret); *Neovison vison* (American mink); *Odocoileus virginianus* (white-tailed deer); *Panthera leo* (lion), *Panthera tigris* (tiger); *Panthera uncia* (snow leopard); *Prionailurus bengalensis* (leopard cat); *Prionailurus viverrinus* (fishing cat); *Mesocricetus auratus* (golden hamster); *Chlorocebus sabaeus* (green monkey) and *Puma concolor* (puma). BLASTx was performed to identify ORF *S* in each variant, and mutations were identified at the same time. Three species (*Mesocricetus auratus, Chlorocebus sabaeus*, and *Puma concolor*) were discarded because they harbored less than three single amino acid mutations. Amino acid mutation data from three additional viruses isolated from mice were retrieved from three studies (Leist et al., 2020; Montagutelli et al., 2021; Sun et al., 2021).

### Estimation of the binding affinity of RBD-ACE2 interaction by molecular docking

We extracted three-dimensional structures of the spike RBD and human ACE2 from the crystal structure (PDB: 6M0J) reported in a previous study (Lan et al., 2020), and those of rodent ACE2 from the predicted models reported in a previous study (Lam et al., 2020). The structure models of the Omicron RBD and the RBD with five mutations (K417N, E484A, Q493R, Q498R, and N501Y) were generated using SWISS-MODEL (Waterhouse et al., 2018), and those of other RBD variants were generated using PyMOL “mutagenesis” (https://pymol.org/). The structure models of the RBD:ACE2 complex were generated by aligning against the reported complex structure of the corresponding species using PyMOL (Lam et al., 2020; Lan et al., 2020).

We performed molecular docking following previous studies (Lam et al., 2020; Rodrigues et al., 2020). Briefly, we refined the three-dimensional models using default refinement protocols, and then estimated the HADDOCK scores for each RBD:ACE2 complex using HADDOCKv2.4 web server (van Zundert et al., 2016). Docking results of each RBD-ACE2 variant pair were clustered, and the average HADDOCK score of the top cluster was reported for the RBD:ACE2 complex.

## ACKNOWLEDGMENTS

We thank Dr. Xionglei He from Sun Yat-sen University and Dr. Mingkun Li from Beijing Institute of Genomics CAS for discussion. We acknowledge the authors and laboratories for generating and submitting the sequences to GISAID Database on which this research is based. The list is detailed in Table S2. This work was supported by grants from the National Natural Science Foundation of China (31922014).

## SUPPLEMENTARY MATERIAL

Supplementary material includes Supplemental Figures S1–4 and Supplemental Tables S1–2.

## AUTHOR CONTRIBUTIONS

W.Q. designed the study; C.W., K.-J.S., W.W., and S.Z. performed data analyses; C.W., K.-J.S., Q.H., and W.Q. wrote the manuscript.

## DATA AVAILABILITY

All scripts used to analyze the data and to generate the figures are available at github (https://github.com/ChangshuoWei/Omicron_origin) and Zenodo (DOI: 10.5281/zenodo.5778199). All data that were used to support the findings of this study are available in the public databases.

## FIGURES

**Fig. S1.**
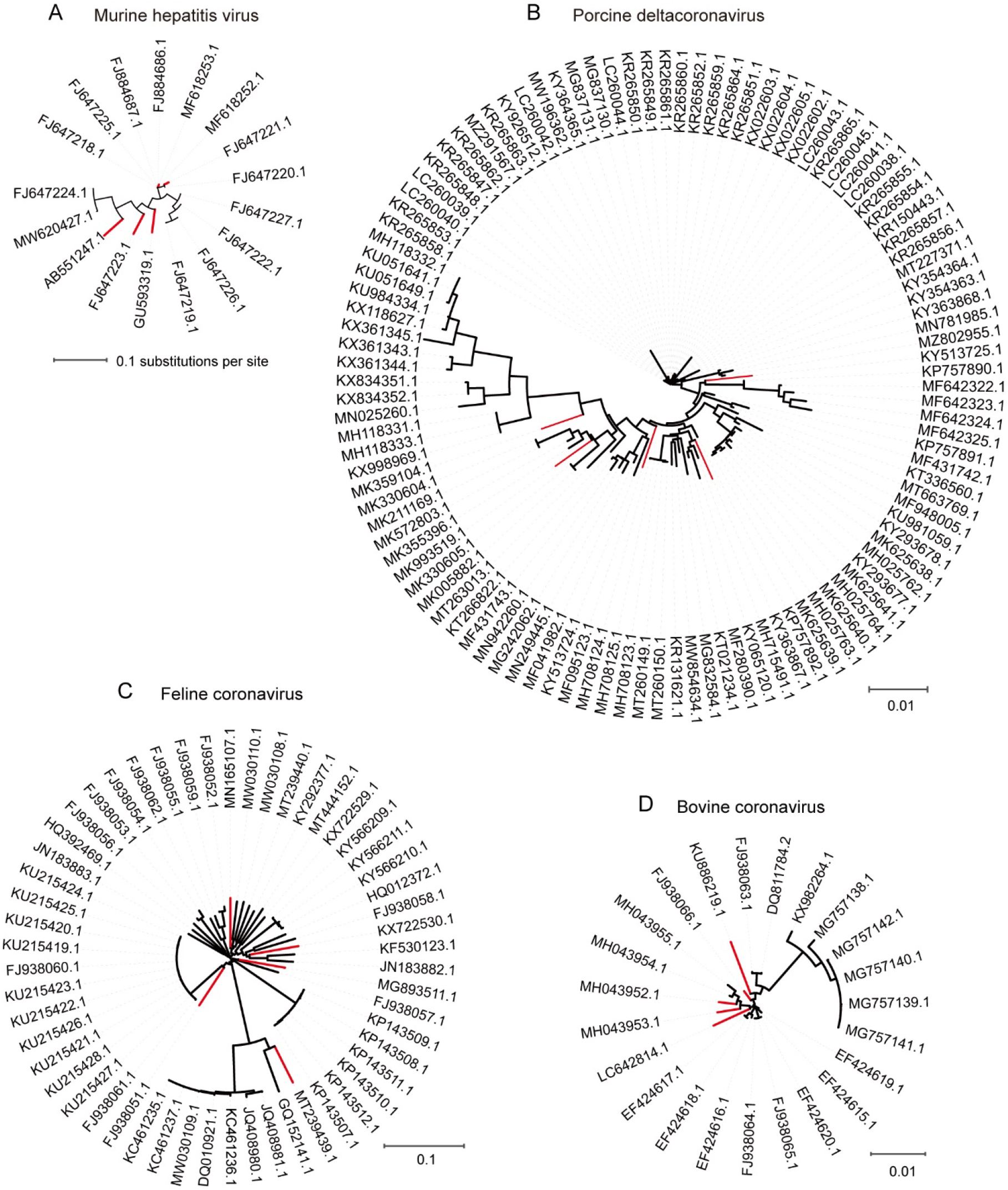
The phylogenetic trees of four coronaviruses species. The five external branches with the largest number of accumulated mutations were colored red, and were used in the principal component analysis.

**Fig. S2.**
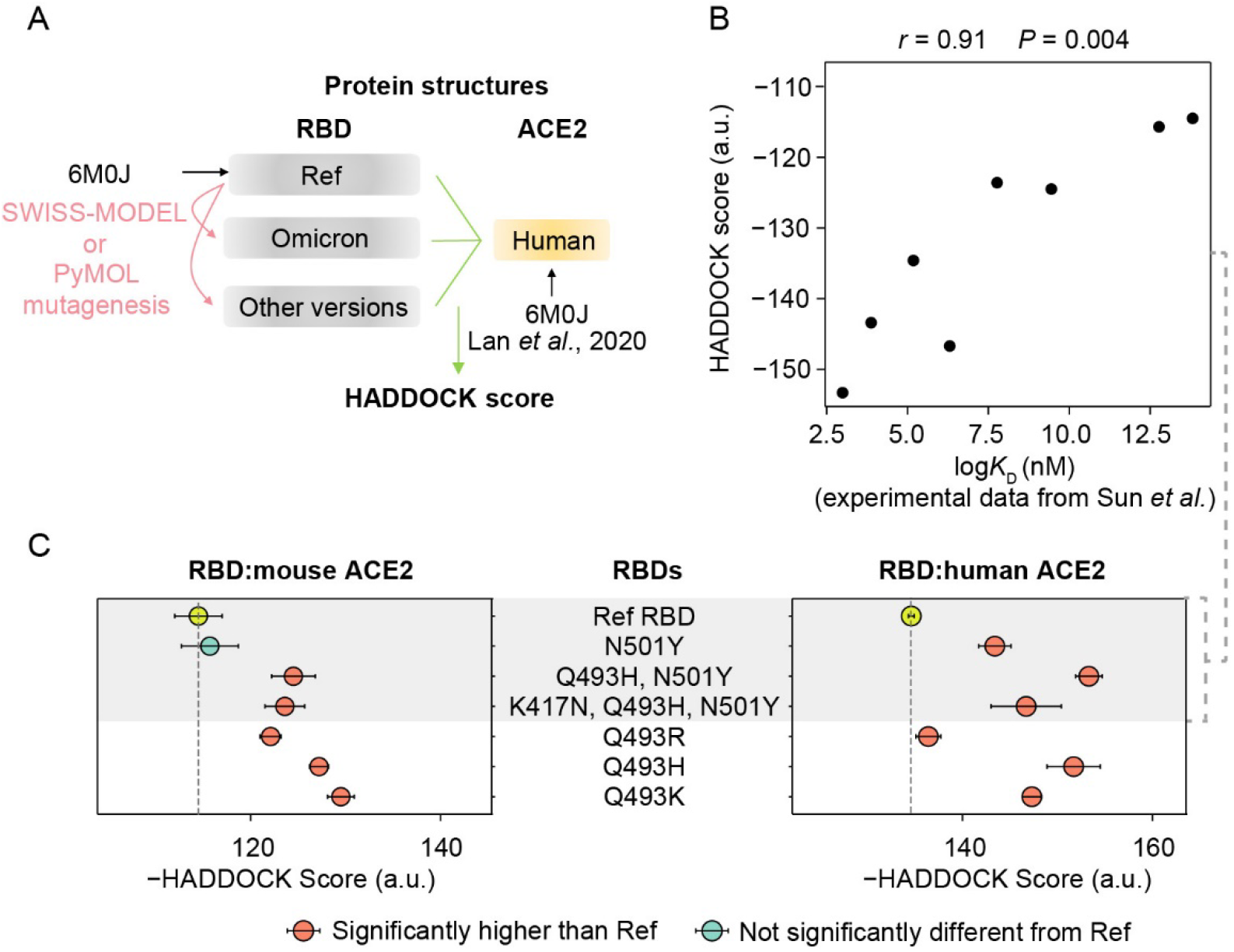
Predicted binding affinities of RBD and ACE2. **A**. A schematic shows the workflow to estimate the HADDOCK scores between RBD variants and human ACE2. **B**. A scatterplot shows the correlation between the estimated HADDOCK score and the *K*_D_ value. The *K*_D_ values were experimentally determined in Sun *et al*. (2021). **C**. The HADDOCK scores for the interactions between RBD variants and mouse/human ACE2. The top four rows were used for drawing the scatterplot in (B). The error bars represent standard errors.

**Fig. S3.**
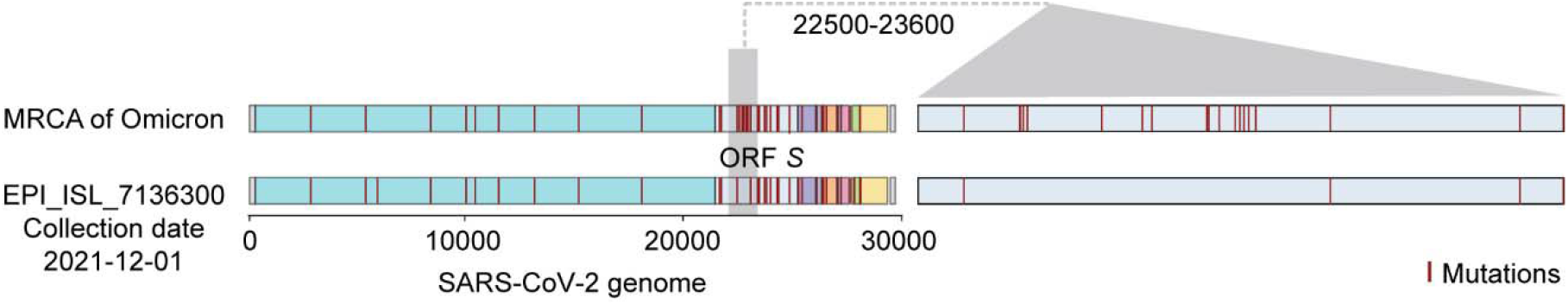
The distribution of mutations in a potential intermediate of the current Omicron variants. Red lines represent the positions of point mutations (relative to the B.1.1 lineage). The genomic regions were colored according to ORFs as in **Fig. 1B**.

**Fig. S4.**
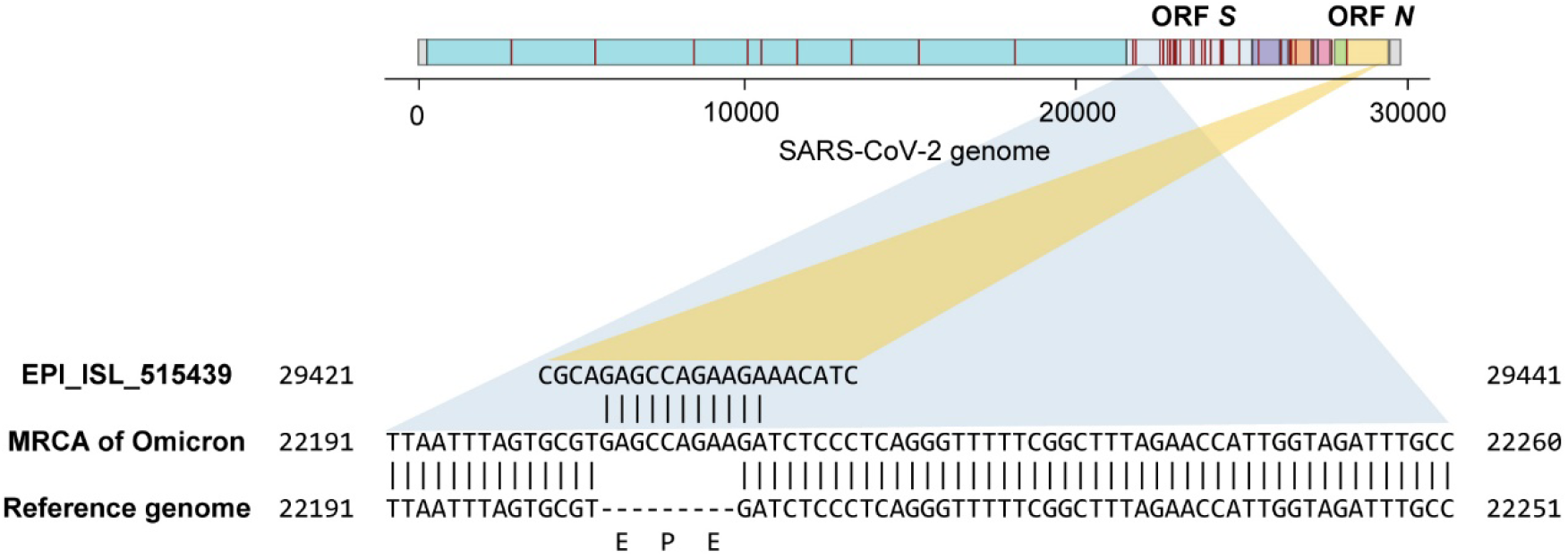
A potential explanation for the “EPE insertion” in the spike protein of Omicron. Red lines represent the positions of point mutations (relative to the B.1.1 lineage). The genomic regions were colored according to ORFs as in **Fig. 1B**. The “EPE insertion” related regions were shown at the nucleotide resolution. The SARS-CoV-2 variant (EPI_ISL_515439) collected by the Nevada State Public Health Laboratory on May 19^th^, 2020 was used as an example to illustrate a potential source sequence of the insertion.

